# A false sense of circularity: hidden variability in RNase R enzymes confounds reliable circRNA validation

**DOI:** 10.64898/2026.10.13.757942

**Authors:** Lotte Ulrix, Jolien De Waele, Jo Vandesompele, Mattias Hofmans, Barbara De Moerloose, Tim Lammens

## Abstract

Circular RNAs (circRNAs) are an emerging class of RNA molecules involved in development and disease. They are generated through back-splicing from pre-mRNA. RNaseR treatment, which degrades linear RNA while preserving circRNAs, is widely used for circRNA validation and enrichment. This short report demonstrated differential performance of two RNaseR enzymes from different suppliers, despite identical experimental conditions, indicating that these enzymes should not be considered interchangeable across suppliers. Importantly, these findings highlight that without demonstrating efficient linear RNA degradation, unchanged circRNA abundance after RNase R treatment is inherently uninformative and may contribute to false-positive circRNA validation or inadequate circRNA enrichment. We therefore highlight that RNaseR digestion should be consistently verified using a linear control, particularly when changing enzyme suppliers

## Manuscript

Although first described in plant viroids fifty years ago and later found among eukaryotic transcripts, circular RNAs (circRNAs) were largely dismissed for decades as splicing noise or artifacts of low abundance [1,2]. This view changed with the advent of high-throughput RNA sequencing and dedicated circRNA detection algorithms, which revealed that these molecules are abundant, conserved, and often have a tissue-specific expression [3-5]. Landmark studies subsequently established that circRNAs may have defined biological functions in development and disease, cementing circRNAs as a distinct and biologically relevant class of noncoding RNA rather than a technical byproduct [5-8].

CircRNAs are formed through back-splicing of precursor mRNA, a non-canonical splicing event in which a downstream splice donor is joined to an upstream splice acceptor. This process results in a covalently closed RNA molecule that lacks a poly(A)tail and free 5’ and 3’ ends, and has a characteristic back-splice junction (BSJ) [9,10]. An empirical challenge in the study of circRNAs is distinguishing genuine circular RNA molecules from co-detected linear transcripts sharing the same exon composition, including trans-spliced linear RNAs, which can generate false-positive back-splice signals in both RNA sequencing and RT-qPCR data. The gold standard for proving circularity exploits a simple biochemical property of circRNAs: RNase R, a 3′→5′ exoribonuclease, degrades linear RNA but cannot digest covalently closed circular molecules lacking free ends [11,12]. Consequently, RNase R treatment of total RNA prior to RT-qPCR or sequencing has become the standard approach to confirm both circularity (by demonstrating stable circRNA abundance alongside depletion of the corresponding linear transcript) and enrich circRNAs before library preparation. It is worth noting that RNase R digestion is not absolute or universally complete since some structured linear RNAs are known to resist digestion, while some long circular RNAs are susceptible to degradation. This illustrates that “RNase R treatment” is not a simple and universally working enzymatic step, but one sensitive to reaction conditions and substrate structure [13,14]. Given how central this step is, the use of a rigorously validated and robust protocol is of paramount importance. Vromman et al. [15] published exactly that: a detailed, validated workflow for RNase R treatment, cleanup, and RT-qPCR-based validation of circRNAs, now widely used as the reference method in the field. In short, total RNA (1 µg) was subjected to RNase R digestion using 1 U RNase R per reaction in a 15 µL reaction volume, with matched buffer-only controls. Samples were incubated for 15 min at 37°C, followed by buffer exchange using Vivacon 500 Hydrosart 10 kDa ultrafiltration columns. Reverse transcription and RT-qPCR were subsequently performed to assess the stability of circRNAs and linear control RNAs following RNase R treatment. Their optimized RNase R workflow achieved near-complete degradation of linear RNAs [ΔCq ≈ 8], while maintaining circRNA stability after digestion [ΔCq ≈ 0.5].

As a first step towards implementing RNases R based RT-qPCR validation of circRNAs in our leukemia research, we sought to reproduce the results obtained by Vromman et al.[15]. Despite strict adherence to the published protocol, including the use of RNA from the same cell line and the same RT-qPCR assay, we failed to degrade linear transcripts: Cq values for the linear control transcripts (mRNA ACTB and TBP) were essentially unchanged before and after RNase R treatment ([ΔCq ≈ 1]), and, unsurprisingly, the circRNA targets (circHIPK3; chr11:33286412-33287511, circSETD3; chr14:99458278-99465813) equally showed minimal change ([ΔCq ≈ 1]). Primer sequences are presented in supplementary Table 1. Without demonstrating efficient degradation of linear control transcripts, stable circRNA Cq values cannot be taken as evidence of RNase R resistance, as they are equally consistent with a failure of the RNase R treatment itself, and thus this pattern is uninformative.

We did however source the RNase R enzyme from Abcam (cat. no. M1228-500), rather than the supplier specified in the original protocol (Epicentre, cat. no. RNR07250). We subsequently evaluated this RNase R enzyme from Epicentre, repeated the identical protocol and reagents in parallel, and obtained results matching the published data: substantial degradation of the linear control mRNAs ([ΔCq ≈ 5]) alongside stable Cq values for the circRNA targets ([ΔCq ≈ 0]) (**Figure 1A**). Additionally, these results were confirmed in an independent leukemia cell line model (KASUMI-1) (**Figure 1B**). The only variable changed between the two experiments was the commercial source of the enzyme. Importantly, both enzymes are derived from E. coli, are supplied at similar concentrations (Abcam: 10 U/µL; Epicentre: 20 U/µL), and have their activity defined identically, with one unit corresponding to the conversion of 1 µg of poly(A) into acid-soluble nucleotides in 10 minutes at 37°C. Thus, no substantial differences were initially expected.

**Figure 1:**
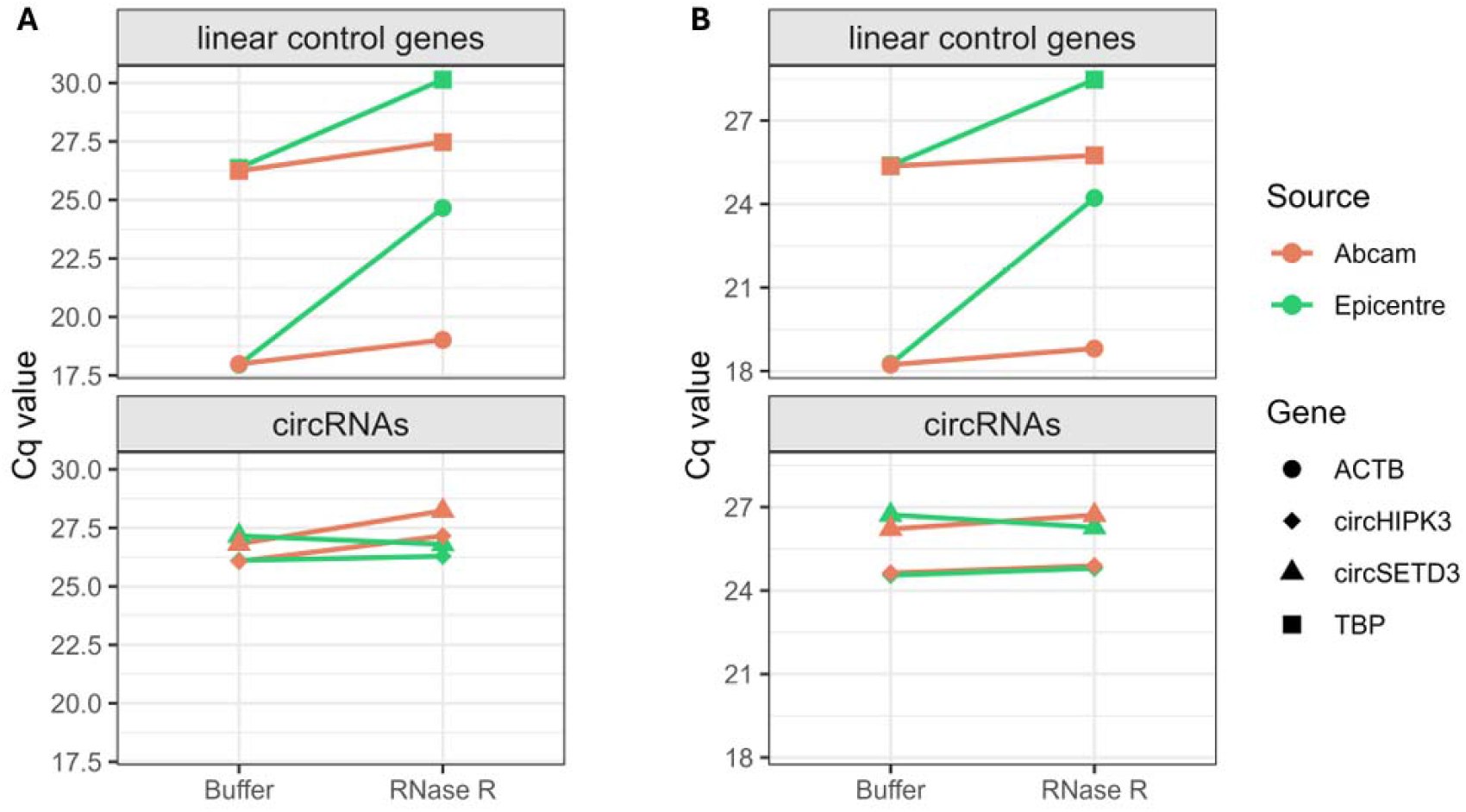
qPCR validation of RNase R digestion using Abcam vs Epicentre enzyme. RNase R and buffer control treatments were performed on total RNA from (A) SW480 and (B) KASUMI-1 in duplicate. Average Cq values for both qPCR replicates are shown for linear control mRNAs (ACTB and TBP) and circRNAs (circHIPK3; chr11:33286412-33287511, circSETD3; chr14:99458278-99465813). A: In SW480 total RNA treated with RNase R from Abcam, an average increase in Cq of 1.13 for linear control mRNAs and 1.26 for circRNAs was observed. In SW480 samples treated with RNase R from Epicentre, an average increase in Cq of 5.24 for linear control mRNAs and 0 for circRNAs was observed. B: In KASUMI-1 total RNA treated with RNase R from Abcam, an average increase in Cq of 0.48 for linear control mRNAs and 0.38 for circRNAs was observed. In KASUMI-1 samples treated with RNase R from Epicentre, an average increase in Cq of 4.52 for linear control mRNAs and 0.1 for circRNAs was observed.

Although admittedly, a small and simple comparison of presumed identical enzymes from two different commercial sources in one experiment, the implication is disproportionate: without an explicit linear RNA digestion control, a failed RNase R digestion can be mistaken for successful validation of circularity. Although the use of a linear RNA control is recommended in the original protocol, it is easily disregarded as a formality rather than a diagnostic checkpoint. Consequentially, a nonfunctional enzyme lot or supplier can silently cause false-positive circRNA validations or, in a sequencing context, fail to enrich for circRNAs while giving the impression of the opposite.

We are not aware of a systematic comparison of commercial RNase R sources for circRNA applications, and we do not claim our two-enzyme comparison to be one. Our purpose here is narrower: to flag that “RNase R” is not an interchangeable commodity reagent across suppliers, and that enzyme source deserves the same scrutiny normally reserved for PCR primers or reference genes. We recommend that any RNase R protocol, regardless of enzyme source, always include and report a parallel linear-RNA digestion control, and that labs switching suppliers or lots first validate digestion efficiency against a known linear transcript before drawing downstream circRNA conclusions.

## Acknowledgments

This work was supported in part by vzw Kinderkankerfonds (T.L.), Stichting Tegen Kanker (B.D.M., 2022-145), Ghent University BOF (J.V., 2021/244) and FWO (J.D.W., 11D4425N).

## Figures

**Supplementary table 1:**
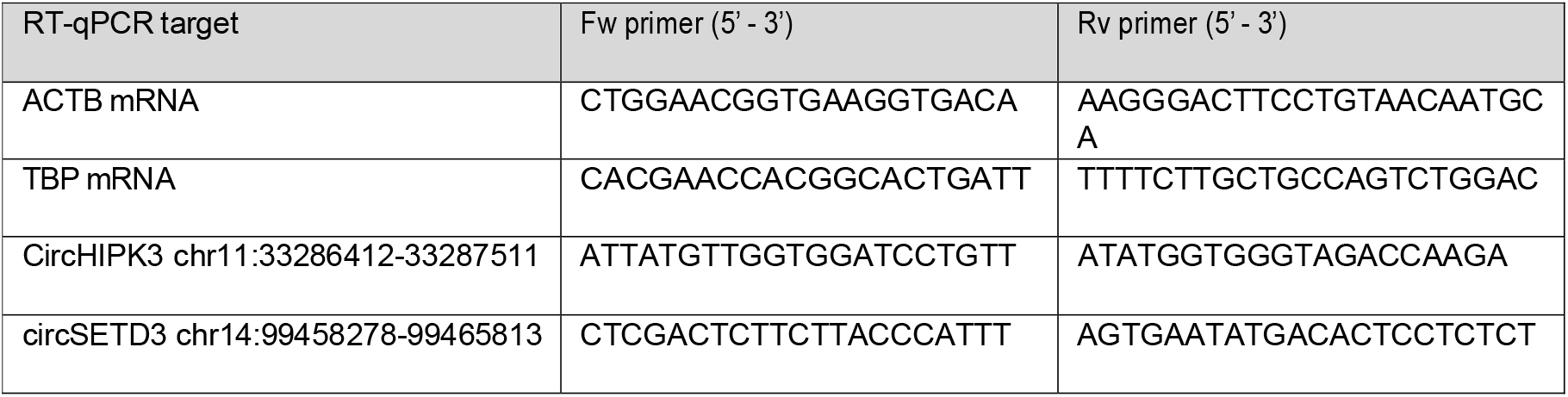
Forward (Fw) and reverse (Rv) primer sequences of linear control genes and circRNAs (Vromman et al. 2021).

## References

[1] Sanger HL, Klotz G, Riesner D, et al. Viroids are single-stranded covalently closed circular RNA molecules existing as highly base-paired rod-like structures. Proc Natl Acad Sci U S A. 1976 Nov;73(11):3852–6.

[2] Wang PL, Bao Y, Yee MC, et al. Circular RNA is expressed across the eukaryotic tree of life. PLoS One. 2014;9(6):e90859.

[3] Jeck WR, Sorrentino JA, Wang K, et al. Circular RNAs are abundant, conserved, and associated with ALU repeats. Rna. 2013 Feb;19(2):141–57.

[4] Xia S, Feng J, Lei L, et al. Comprehensive characterization of tissue-specific circular RNAs in the human and mouse genomes. Brief Bioinform. 2017 Nov 1;18(6):984–992.

[5] Memczak S, Jens M, Elefsinioti A, et al. Circular RNAs are a large class of animal RNAs with regulatory potency. Nature. 2013 Mar 21;495(7441):333–8.

[6] Hansen TB, Jensen TI, Clausen BH, et al. Natural RNA circles function as efficient microRNA sponges. Nature. 2013 Mar 21;495(7441):384–8.

[7] Chen L, Shi J, Wu Y, et al. CircRNA CDR1as promotes hepatoblastoma proliferation and stemness by acting as a miR-7-5p sponge to upregulate KLF4 expression. Aging (Albany NY). 2020 Oct 14;12(19):19233–19253.

[8] Seeler S, Andersen MS, Sztanka-Toth T, et al. A Circular RNA Expressed from the FAT3 Locus Regulates Neural Development. Mol Neurobiol. 2023 Jun;60(6):3239–3260.

[9] Salzman J, Gawad C, Wang PL, et al. Circular RNAs are the predominant transcript isoform from hundreds of human genes in diverse cell types. PLoS One. 2012;7(2):e30733.

[10] Ashwal-Fluss R, Meyer M, Pamudurti NR, et al. circRNA biogenesis competes with pre-mRNA splicing. Mol Cell. 2014 Oct 2;56(1):55–66.

[11] Suzuki H, Zuo Y, Wang J, et al. Characterization of RNase R-digested cellular RNA source that consists of lariat and circular RNAs from pre-mRNA splicing. Nucleic Acids Res. 2006 May 8;34(8):e63.

[12] Chu LY, Hsieh TJ, Golzarroshan B, et al. Structural insights into RNA unwinding and degradation by RNase R. Nucleic Acids Res. 2017 Nov 16;45(20):12015–12024.

[13] Xiao MS, Wilusz JE. An improved method for circular RNA purification using RNase R that efficiently removes linear RNAs containing G-quadruplexes or structured 3’ ends. Nucleic Acids Res. 2019 Sep 19;47(16):8755–8769.

[14] Rahimi K, Venø MT, Dupont DM, et al. Nanopore sequencing of brain-derived full-length circRNAs reveals circRNA-specific exon usage, intron retention and microexons. Nat Commun. 2021 Aug 10;12(1):4825.

[15] Vromman M, Yigit N, Verniers K, et al. Validation of Circular RNAs Using RT-qPCR After Effective Removal of Linear RNAs by Ribonuclease R. Curr Protoc. 2021 Jul;1(7):e181.

