## Supplementary Table 1 for "A false sense of circularity: hidden variability in RNase R enzymes confounds reliable circRNA validation"

Supplementary table 1: Forward (Fw) and reverse (Rv) primer sequences of linear control genes and circRNAs (Vromman et al. 2021).

| RT-qPCR target | Fw primer (5’ - 3’) | Rv primer (5’ - 3’) |
| --- | --- | --- |
| ACTB mRNA | CTGGAACGGTGAAGGTGACA | AAGGGACTTCCTGTAACAATGCA |
| TBP mRNA | CACGAACCACGGCACTGATT | TTTTCTTGCTGCCAGTCTGGAC |
| CircHIPK3 chr11:33286412-33287511 | ATTATGTTGGTGGATCCTGTT | ATATGGTGGGTAGACCAAGA |
| circSETD3 chr14:99458278-99465813 | CTCGACTCTTCTTACCCATTT | AGTGAATATGACACTCCTCTCT |
